# The Monarch Initiative: An integrative data and analytic platform connecting phenotypes to genotypes across species

**DOI:** 10.1101/055756

**Authors:** Christopher J Mungall, Julie A McMurry, Sebastian Köhler, James P. Balhoff, Charles Borromeo, Matthew Brush, Seth Carbon, Tom Conlin, Nathan Dunn, Mark Engelstad, Erin Foster, JP Gourdine, Julius O.B. Jacobsen, Daniel Keith, Bryan Laraway, Suzanna E. Lewis, Jeremy Nguyen Xuan, Kent Shefchek, Nicole Vasilevsky, Zhou Yuan, Nicole Washington, Harry Hochheiser, Tudor Groza, Damian Smedley, Peter N. Robinson, Melissa A Haendel

**Affiliations:** Environmental Genomics and Systems Biology, Lawrence Berkeley National Laboratory, Berkeley, CA, USA; Department of Medical Informatics and Clinical Epidemiology and OHSU Library, Oregon Health & Science University, Portland, USA; Institute for Medical Genetics and Human Genetics, Charité-Universitätsmedizin Berlin, Augustenburger Platz 1, 13353 Berlin, Germany; RTI International, Research Triangle Park, NC, USA; Department of Biomedical Informatics, University of Pittsburgh, Pittsburgh, PA; Wellcome Trust Sanger Institute, Hinxton, Cambridge, CB10 1SA, UK; Kinghorn Centre for Clinical Genomics, Garvan Institute of Medical Research, Darlinghurst, NSW 2010, Australia; William Harvey Research Institute, Barts & The London School of Medicine & Dentistry, Queen Mary University of London, Charterhouse Square, London, EC1M 6BQ, UK

## Abstract

The principles of genetics apply across the whole tree of life: on a cellular level, we share mechanisms with species from which we diverged millions or even billions of years ago. We can exploit this common ancestry at the level of sequences, but also in terms of observable outcomes (phenotypes), to learn more about health and disease for humans and all other species. Applying the range of available knowledge to solve challenging disease problems requires unified data relating genomics, phenotypes, and disease; it also requires computational tools that leverage these multimodal data to inform interpretations by geneticists and to suggest experiments. However, the distribution and heterogeneity of databases is a major impediment: databases tend to focus either on a single data type across species, or on single species across data types. Although each database provides rich, high-quality information, no single one provides unified data that is comprehensive across species, biological scales, and data types. Without a big-picture view of the data, many questions in genetics are difficult or impossible to answer. The Monarch Initiative (https://monarchinitiative.org) is an international consortium dedicated to providing computational tools that leverage a computational representation of phenotypic data for genotype-phenotype analysis, genomic diagnostics, and precision medicine on the basis of a large-scale platform of multimodal data that is deeply integrated across species and covering broad areas of disease.

## Introduction

A fundamental axiom of biology is that phenotypic manifestations of an organism are due to interaction between genotype and environmental factors over time. In the rapidly advancing era of genomic medicine, a critical challenge is to identify the genetic etiologies of Mendelian disease, cancer, and common and complex diseases, and translate basic science to better treatments. Currently, available human data associates less than approximately 51% of known human coding genes with phenotype data (based on OMIM (1), ClinVar (2), Orphanet (3), CTD (4), and the GWAS catalog (5)). See Table 1 for a list of database abbreviations. This coverage can be extended to approximately 89% if phenotypic information from orthologous genes from five of the most well-studied model organisms is included (**Figure 1**).

**Figure 1.**
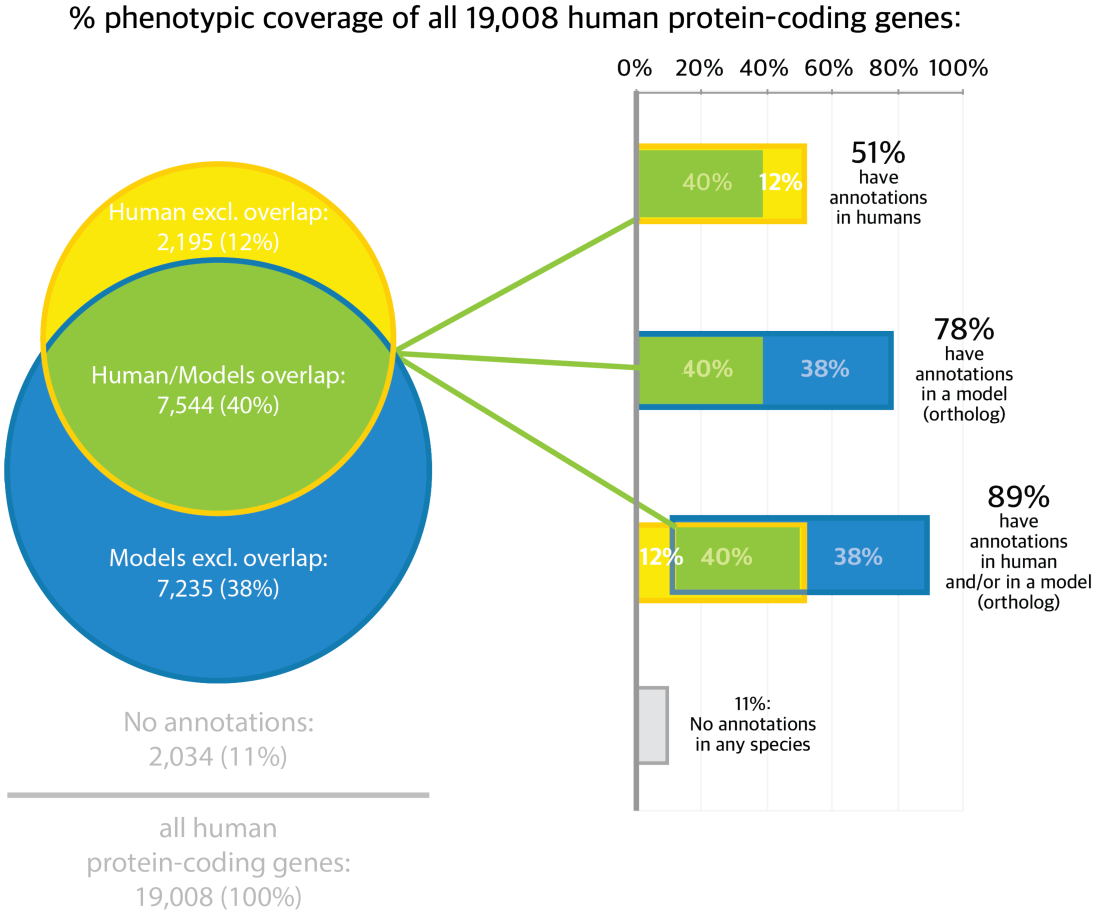
*The phenotype annotation coverage of human coding genes. Yellow bars show that 51% of those genes have at least one phenotype association reported in humans (HPO annotations of OMIM, ClinVar, Orphanet, CTD, and GWAS). The blue bars show that 58% of human coding genes have orthologs with causal phenotypic associations reported in at least one non-human model (MGI, Wormbase, Flybase, and ZFIN). The green bars show that 40% of human coding genes have annotations both in human and in non-human orthologs. There are phenotypic associations from humans and/or non-human orthologs that cover 89% of human coding genes.*

Similarly, of the 72% of the 3,230 genes in ExAC with ‘near-complete depletion of predicted protein-truncating variants have no currently established human disease phenotype’ (6), where 88% of these genes without a human phenotype have a phenotype in a non-human organism. However, leveraging these model data for computational use is non-trivial primarily because the relationships between gene and disease (7) and between model system and disease phenotypes (8) are not straightforward.

In recent years, there has been a growth in the number of genotype-phenotype databases available, covering a diversity of domain areas for human, model organisms, and veterinary species. While providing quality inventories of the relevant species and phenotypic data types, most resources are limited to a single species or limit cross-species comparison to direct assertions (e.g. Organism X is a model of Disease Y) or based upon orthology relations (e.g. organism Y’ is a model of Disease Y due to Y and Y’ being orthologs). While great strides have been made in text-based search engines, phenotype data remains difficult to search and use computationally due to its complexity and in the use of different phenotype standards and terminologies. Such barriers have made linking and integration with the precision and richness needed for mechanistic discovery across species a significant challenge (9). A newer method to aid identifying models of disease and to discovery underlying mechanisms is to utilize ontologies to describe the set of phenotypes that present for a given genotype or disease, what we call a ‘phenotypic profile’. A phenotypic profile is the subject of non-exact matching within and across species using ontology integration and semantic similarity algorithms (10)(11) in software applications such as Exomiser (12) and Genomiser (13), and this approach has been shown assist disease diagnosis (14–16). The Monarch Initiative uses an ontology-based strategy to deeply integrate genotype-phenotype data from many species and sources, thereby enabling computational interrogation of disease models and complex relationships between genotype and phenotype to be revealed. The Monarch Initiative name was chosen because it is a community effort to create paths for diverse data to be put to use for disease discovery, not unlike the navigation routes that a monarch butterfly would take.

## Data Architecture

The overall data architecture for Monarch is shown in **Figure 2**. The bulk of the data integration is carried out using our Data Ingest Pipeline (Dipper) tool (https://github.com/monarch-initiative/dipper), which maps a variety of external data sources and databases to RDF (Resource Description Framework) graphs. RDF provides a flexible way of modeling a variety of complex datatypes, and allows entities from different databases to be connected via common instance or class URIs (Uniform Resource Indicators). We use relationship types from the Relation Ontology (RO; <u>https://github.com/oborel/obo-relations)</u> (17) and other vocabularies to connect entities together, along with a number of Open Biological Ontologies (18) (OBOs) to classify these entities. For example, a mouse genotype can be related to a phenotype using the *has_phenotype* relation (RO:0002200), with the genotype classified using a term from the Genotype Ontology (GENO) (19), and the phenotype classified using the Mammalian Phenotype Ontology (MP) (20). We use the Open Biomedical Annotations (OBAN; https://github.com/EBISPOT/OBAN) vocabulary to associate evidence and provenance metadata with each edge, using the Evidence and Conclusions Ontology (ECO) for types of evidence (21). The graphs produced by Dipper are available as a standalone resource in RDF/turtle format at <u>http://data.monarchinitiative.org/ttl</u>.

**Figure 2.**
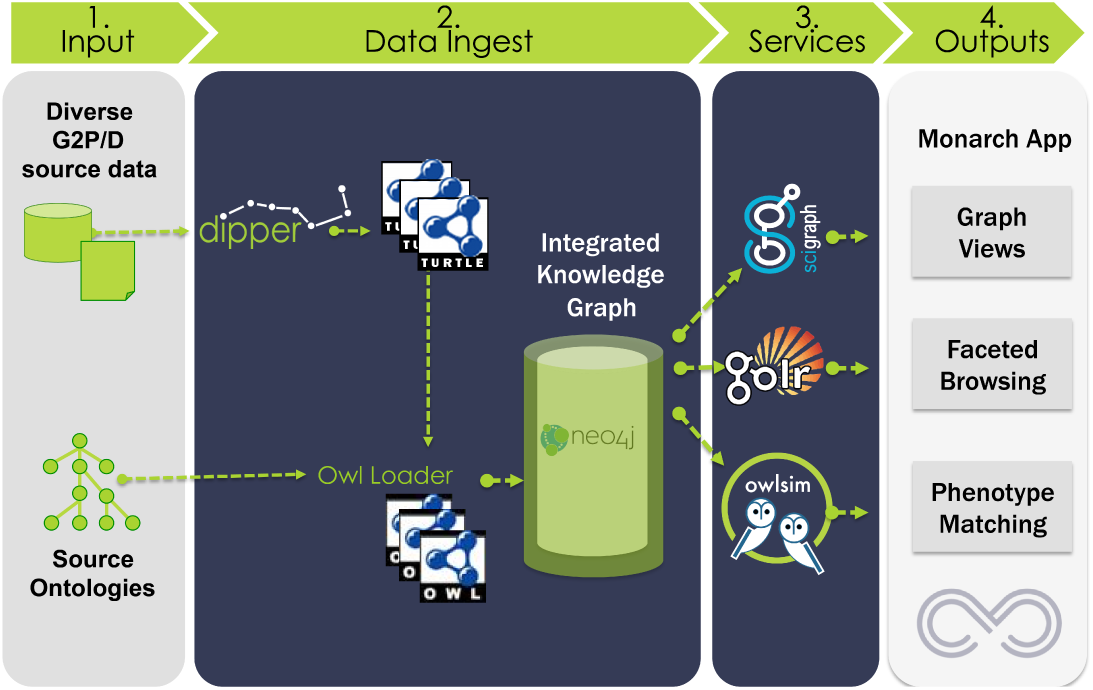
*Monarch Data Architecture. Structured and unstructured data sources are loaded into SciGraph via Dipper. Ontologies are also loaded into SciGraph, resulting in a combined knowledge and data graph. Data is disseminated via SciGraph Services, an ontology-enhanced Solr instance called GOlr, and to the OwlSim semantic similarity software. Monarch applications and end users access the services for graph querying, application population, and phenotype matching.*

We also import a number of external and in-house ontologies, for data description and data integration. As these ontologies are all available from the OBO Library in Web Ontology Language (OWL), no additional transformation is necessary. The combined corpus of graphs ingested using Dipper and from ontologies is referred to as the **Monarch Knowledge Graph**.

The data integrated in Monarch wide range of sources, including human clinical knowledge sources as well as genetic and genomic resources covering organismal biology. The list of data sources and ontologies integrated is shown in **Figure 3**, with a species distribution illustrated in **Figure 4**. The knowledge graph is loaded into an instance of a SciGraph database (<u>https://github.com/SciGraph/SciGraph/</u>), which embeds and extends a Neo4J database, allowing for complex queries and ontology-aware data processing and Named Entity Recognition. We provide two public endpoints for client software to query these services: <u>https://scigraph-ontology.monarchinitiative.org/scigraph/docs</u>(for ontology access) and <u>https://scigraph-data.monarchinitiative.org/scigraph/docs</u>(for ontology plus data access).

**Figure 3.**
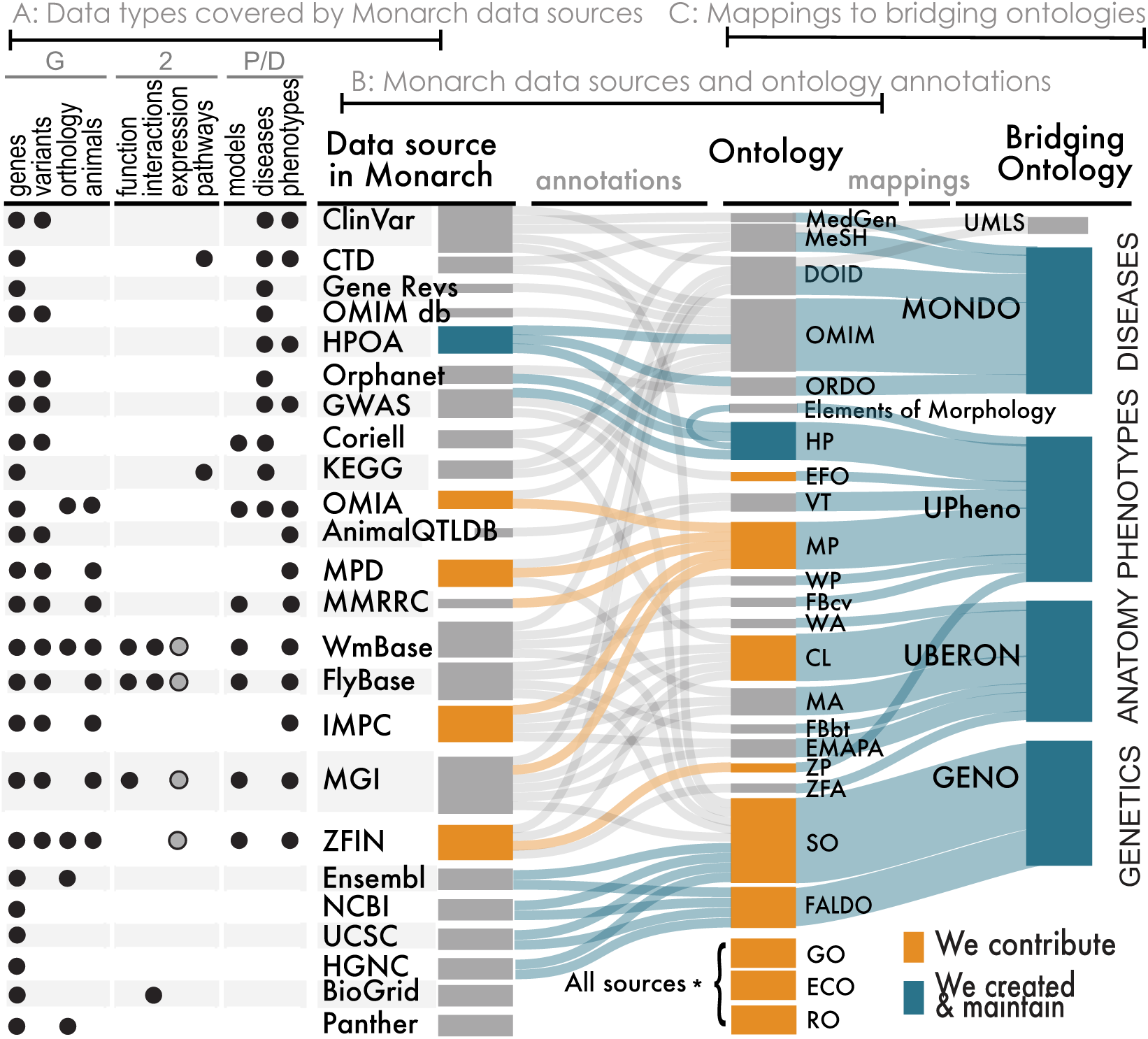
*Data types, sources, and the ontologies used for their integration into the Monarch knowledge graph. Each data source uses or is mapped to a suite of different ontologies or vocabularies. These are in turn integrated into bridging ontologies for Genetics (GENO), Anatomy (Uberon/CL), Phenotypes (UPheno), and Diseases (MonDO).*

**Figure4.**
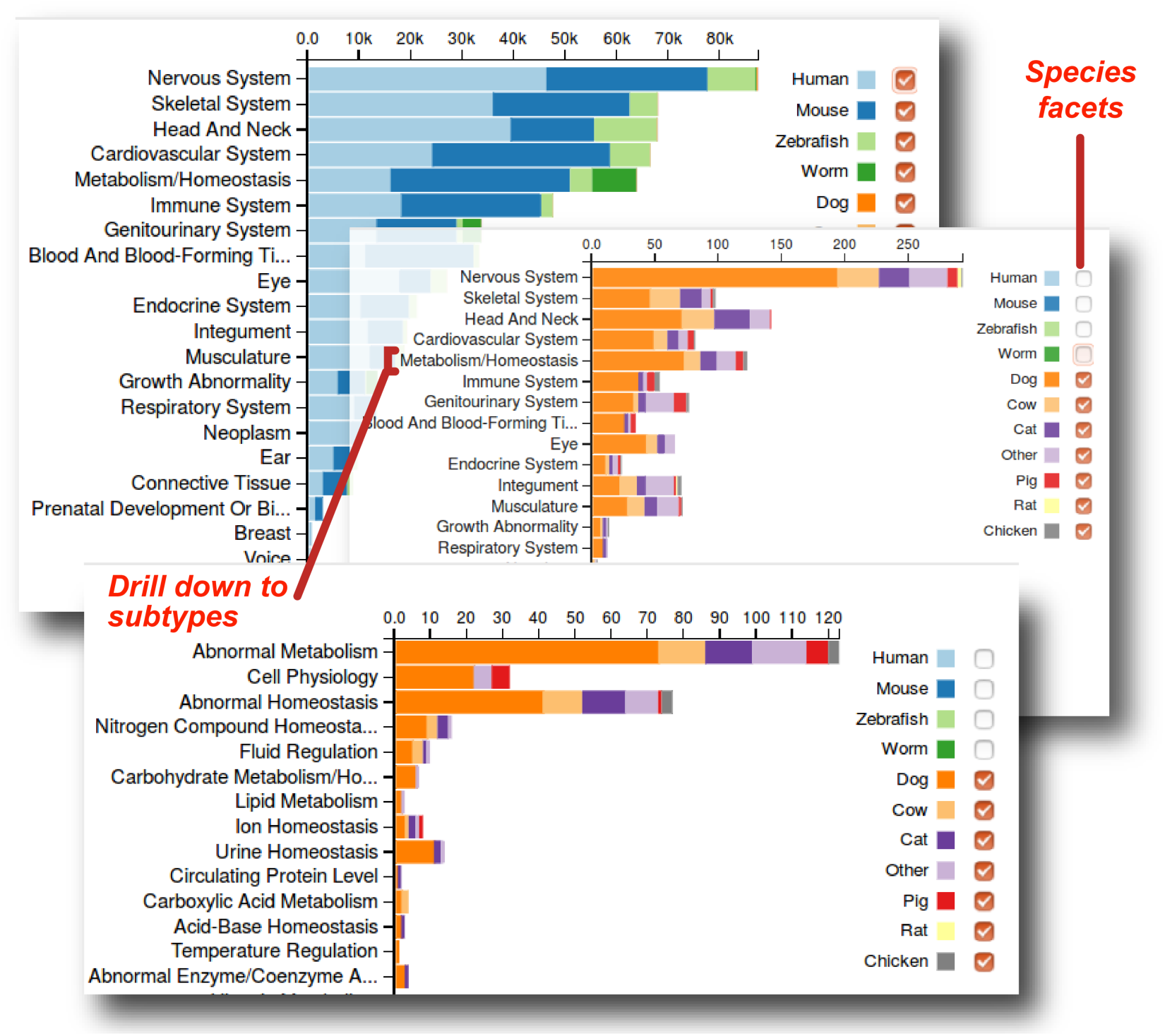
*Distribution of phenotypic annotations across species in Monarch, broken down by the top levelsofthe phenotype ontology. The graph can be interactively explored at <u>https://monarchinitiative.org/phenotype/</u>. Note that annotations are currently dominated by human, mouse, zebrafish and C elegans (top panel); the chartisfacetedallowing individual species to be switched on and off to see contributions for less data-rich species such as veterinary animals and monkeys(middle panel). Clicking on a given phenotype text allows drilling down to its subtypes (lower panel).*

These SciGraph instances provide powerful graph querying capabilities over the complete knowledge graph. Many of the common query patterns are executed in advance and stored in an Apache Solr index, making use of the Gene Ontology ‘GOlr’ indexing strategy, allowing for fast queries of ontology-indexed associations.

Finally, we also load a subset of the graph into an OwlSim instance, which provides phenotype matching services as well as the ability to perform fuzzy phenotype searches based on a phenotype profile. We also provide phenotype matching services via the Global Alliance for Genomes and Health (GA4GH) Matchmaker Exchange (MME) API MME (22), available at <u>https://mme.monarchinitiative.org</u>.

Many of the data sources we integrate make use of their own terminologies and ontologies. We aggregate these into a unified ontology (<u>https://github.com/monarch-initiative/monarch-ontology/</u>) and make use of bridging ontologies and our curated integrative ontologies to connect these together. In particular:

- The Uber-anatomy ontology (Uberon) bridges species-specific and clinical anatomical and tissue ontologies (23)
- The unified phenotype ontology bridges model organism and human phenotype ontologies and terminologies, using techniques described in (24)(25)
- The Monarch Merged Disease Ontology (MonDO) uses a Bayes ontology merging algorithm (26) to integrate multiple human disease resources into a single ontology, and additionally includes animal diseases from OMIA.
- The Genotype Ontology (GENO) (19) defines genotypic elements and bridges the Sequence Ontology (SO) (27) and FALDO (28). GENO allows the propagation of phenotypes that are annotated differently in different sources to genotypic elements.

### Entity resolution and unification

One of the many challenges faced when integrating bioinformatics resources is the presence of the same entity in multiple databases, designated by different identifiers (29). This problem is compounded by the different ways the same identifier can be written, using different prefixes or no prefix at all. Taking a Monarch page for a single gene, for example ‘fibrinogen gamma chain’, *FGG*, (<u>https://monarchinitiative.org/gene/NCBIGene:2266</u>). Monarch has integrated data from a variety of human, model organism, and other biomedical sources such as OMIM (1), Orphanet (3), ClinVar (2), HPO (30), KEGG (31), CTD (4), MyGene (32), BioGrid (33), and via orthology in PantherDB (34) we also incorporate *Fgg* gene data from ZFIN (35) and from MGI (36). No two of these sources represents the identifier for *FGG* in precisely the same way. As part of our data ingest process, we normalize all identifiers using a curated set of database prefixes. These have a defined mapping to an http URI. These curated prefixes have been deposited in the Prefix Commons (<u>https://github.com/prefixcommons)</u>, which similarly contains identifier prefixes used within the Gene Ontology (37) and Bio2RDF (38).

In post-processing equivalent identifiers, we perform clique-merging (<u>https://github.com/SciGraph/SciGraph/wiki/Post-processors</u>). We take all edges labeled with either the owl:sameAs or owl:equivalentClasses property and calculate equivalence cliques, based on the symmetric and transitive nature of these properties. We then merge these cliques together, taking a designated ‘clique leader’ (for instance, NCBI for genes) and mapping all edges in the monarch graph such that they point to a clique leader.

### In-house curation

In addition to ingest of external sources and ontologies, we perform in-house data and ontology curation. For curation of ontology-based genotype-phenotype associations (including disease-phenotypic profiles), we are transitioning to the WebPhenote platform (<u>http://create.monarchinitiative.org</u>), which allows a variety of disease entities to be connected to phenotypic descriptors. We also make use of text mining to create seed disease-phenotype associations using the Bio-Lark toolkit (39), which are then manually curated. Most recently we have performed a large-scale annotation of PubMed to extract common disease-phenotype associations (40). Most of the in-house curation work involves making smaller resources with free text description of phenotypic information computable, for example, the Online Mendelian Inheritance in Animals (OMIA) resource, with whom we have been collaborating to support this curation (41).

### Quality control

External resources and datasets that are incorporated into Monarch are evaluated before incorporation into the Dipper pipeline – we primarily integrate high-quality curated resources. For all ontologies we bring in, we apply automated reasoning to detect inconsistencies between different ontologies. For each release, we perform high-level checks on each integrated resource to ensure no errors in the extraction process occurred, but we do not perform in-depth curation checks of integrated resources. Each release happens once every one to two months.

In order to measure annotation richness, we have also created an annotation sufficiency meter web service (42) available at https://monarchinitiative.org/page/services; this service determines whether a given phenotype profile for any organism is sufficiently broad and deep to be of diagnostic utility. The sufficiency score can be displayed as a five star scale as in PhenoTips (43) and in the Monarch web portal (see below) to aid curation or data entry, and can also be used to suggest additional phenotypic assays to be performed – whether in a patient or in a model organism.

## Monarch Web Portal

The Monarch portal is designed with a number of different use cases in mind, including:

- A researcher interested in a human gene, its phenotypes, and the phenotypes of orthologs in model organisms and other species
- Patients or researchers interested in a particular disease or phenotype (or groups of these), together with information on all implicated genes
- A clinical scenario in which a patient has an undiagnosed disease showing a spectrum of phenotypes, with no definitive candidate gene demonstrated by sequencing; in this scenario the clinician wishes to search for either known diseases that have a similar presentation, or model organism genes that demonstrate homologous phenotypes when the gene is perturbed
- Researcher looking for diseases that have similar phenotypic feature to a newly identified model organism mutant identified in a screen
- Researchers or clinicians who need to identify potentially informative phenotyping assays for differential diagnosis or to identify candidate genes

### Features

#### Integrated information on entities of interest

We provide overview pages for entities such as genes, diseases, phenotypes, genotypes, variants, and publications. Each page highlights the provenance of the data from the diverse clinical, model organism, and non-model organism sources. These pages can be found either via search (see below) or through an entity resolver. For example, the URL <u>https://monarchinitiative.org/OMIM:266510</u> will redirect to a page about the disease ‘Peroxisome biogenesis disorder type 3B’ from the OMIM resource, showing its relationships to other content within the Monarch knowledge graph, such as phenotypes and genes associated with the disease. We make use of MonDO (the Monarch merged disease ontology (26)) to group similar diseases together. **Figure 5** shows an example page for Marfan syndrome with related phenotype, gene, model and variant data.

**Figure 5.**
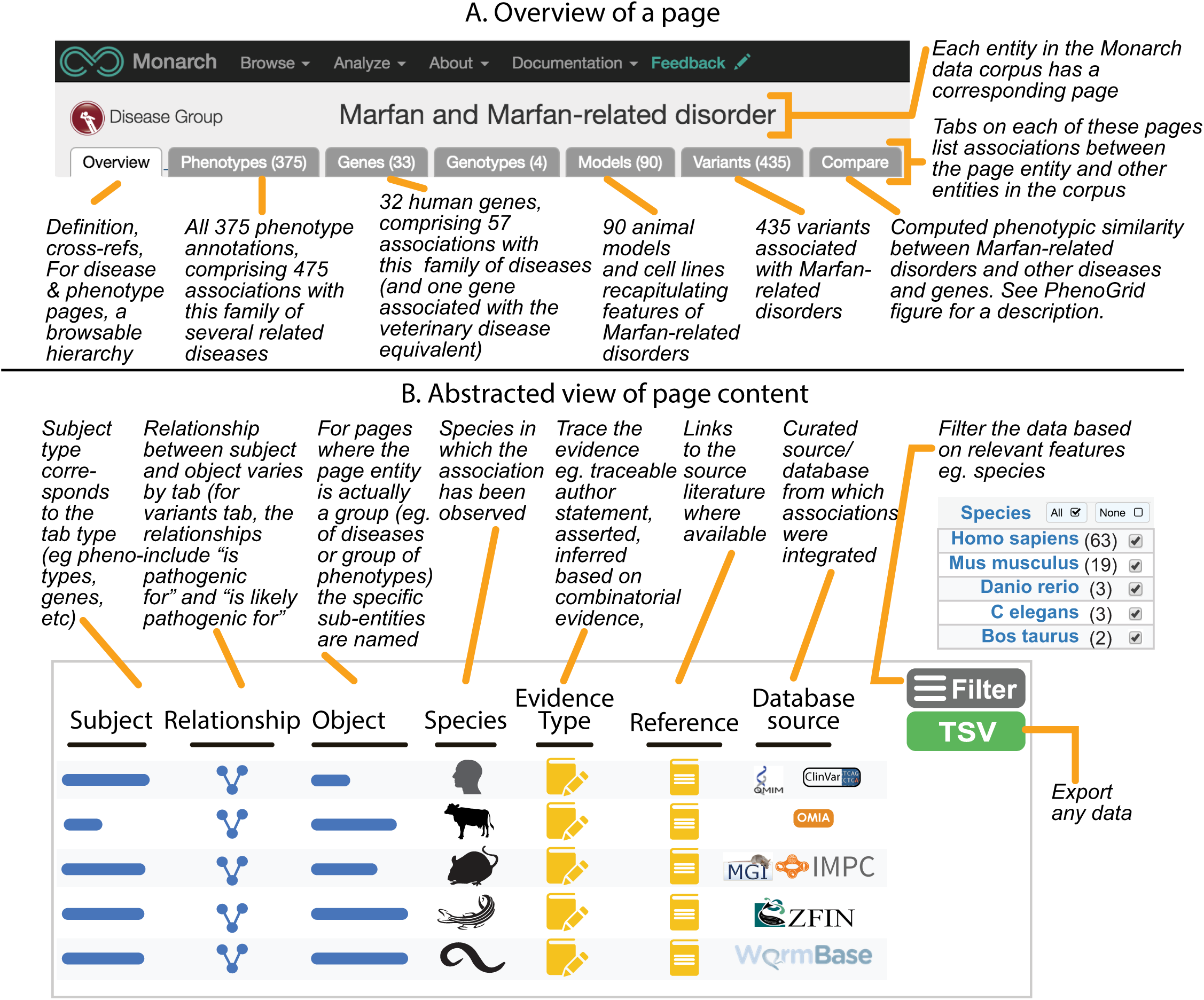
*Annotated Monarch webpage for Marfan and Marfan Related syndrome. This group of syndromic diseases has a number of different associations spanning multiple entity types – disease phenotypes, implicated human genes, variants and animal models and other model systems. An abstraction of the contents and features of the tabs is shown in the lower panel. Actual contents of the tabs are best viewed in the context of the web app at https://monarchinitiative.org/DOID:14323.*

#### Basic Search

The portal provides different means of searching over integrated content. In cases where a user is interested in a specific disease, gene, phenotype etc., these can usually be found via autocomplete. Site-wide synonym-aware text search can also be used to find pages of interest. Because the knowledgebase combines information from multiple species, entities such as genes often have ambiguous symbols. We provide species information to help disambiguate in a search.

#### Search by phenotype profile

One of the most innovative features of Monarch is the ability to query within and across species to look for diseases or organisms that share a set of similar but non-exact set of phenotypes (phenotypic profile). This feature uses a semantic similarity algorithm available from the OWLsim package (<u>http://owlsim.org</u>). Users can launch searches against specific targets: organisms, sets of named gene models, or against all models and diseases available in the Monarch repository. The Monarch Analyze Phenotypes interface (<u>https://monarchinitiative.org/analyze/phenotypes</u>) allows the user to build up a ‘cart’ of phenotypes, and then perform a comparison against phenotypes related to genes and diseases, Results are ranked according to closeness of match, partitioned by species, and are displayed as both a list and in the Phenogrid widget (**Figure 6**).

**Figure 6.**
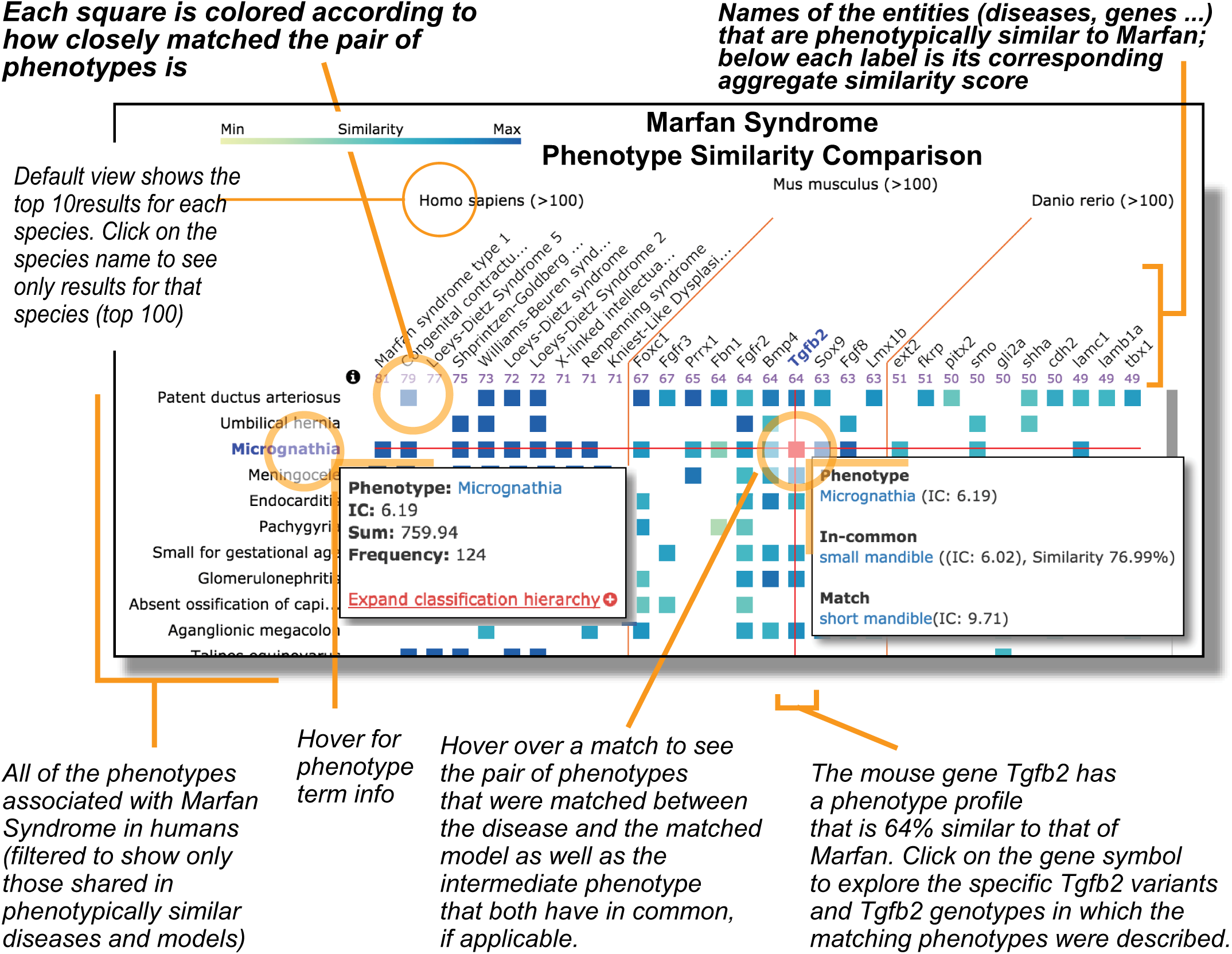
*Partial screenshot of PhenoGrid showing Marfan syndrome. PhenoGrid shows input phenotypes in rows, models in columns, and cell contents color-coded with greater saturation indicating greater similarity. Disease phenotypes are shown as rows, and phenotypically matching human diseases and model organism genes are shown as columns – the saturation of a cell correlates with strength if phenotypic match. Mouse-over tooltips highlight diseases associated with a selected phenotype (or vice-versa), or details (including similarity scores) of any match between a phenotype and a model. User controls support the selection of alternative sort orders, similarity metrics, and displayed organism(s) (mouse, human, zebrafish, or the 10 most similar models for each). Here we see all diseases or genes that exhibit ‘Hypoplasia of the mandible’ with the matching mouse gene Tfgb2. Actual PhenoGrid data is best viewed in the context of the web app at https://monarchinitiative.org/Orphanet:284993#compare.* *Note matches do not need to be exact – here the mouse phenotype of ‘small mandible’ (Mouse Phenotype Ontology) has a high scoring match to “micrognathia” (Human Phenotype Ontology) based on the fact that both phenotypes are related to “small mandible” (Mouse Phenotype Ontology). Advanced PhenoGrid features (not displayed) include the ability to alter the scoring and sorting methods, as well as zoomed-out map-style navigation.*

#### Phenogrid

Given a set of input phenotypes, as associated with a patient or a disease, Monarch phenotypic profile similarity calculations can generate results involving hundreds of diseases and models. The PhenoGrid visualization widget (**Figure 6**) provides an overview of these similarity results, implemented using the D3 javascript library (44). Phenotypes and models are frequently too numerous to fit on the initial display; thus scrolling, dragging, and filtering have been implemented. PhenoGrid is available as an open-source widget suitable for integration in third-party web sites, such as for model organism databases as done in the International Mouse Phenotyping Consortium (IMPC) or clinical comparison tools. Download and installation instructions are available on the Monarch Initiative web site.

#### Text Annotation

The Monarch annotation service allows a user to enter free text (for example, a paper abstract or a clinical narrative) and perform an automated annotation on this text, with entities in the text marked up with terms from the Monarch knowledge graph, such as genes, diseases, and phenotypes. Once the text is marked up, the user has the option of turning the recognized phenotype terms into a phenotype profile, and performing a profile search, or to link to any of the entity pages identified in the annotation. This tool is also available via services.

#### Inferring causative variants

The Exomiser (45) and more recently, Genomiser tools (46) make use of the Monarch platform and phenotype matching algorithms to rank putative causative variants using a combined variant and phenotype score. These tools have been used to diagnose patients as part of the NIH Undiagnosed Diseases Project (14) and are the first examples of using model organism phenotype data to aid rare disease diagnostics.

## Discussion

The Monarch Initiative provides a system to organize and harmonize the heterogeneous genotype-phenotype data found across clinical and model and non-model organism resources (such as veterinary species), creating a unified overview of this rich landscape of data sources. Some of the challenges we have had to address are that each resource shares data via different mechanisms and uses a different data model. It is particularly important to note that each organism annotates phenotypic data to different aspects of the genotype – one resource might be to a gene, another an allele, another to a set of alleles, a full genotype or a SNP. This not only makes data integration difficult, but it also means that computation over the genotype-phenotype associations must be done with care. Similar issues at MGI have been described (47). In addition, since most anatomy, phenotype, and disease ontologies describe the biology of one species, it has traditionally been quite difficult to ‘map’ across species. Some examples are the Human Phenotype Ontology (HPO) (30) and the Mouse Anatomy Ontology (48). Monarch uses four species-neutral ontologies that unify their species-specific counterparts (as shown in Figure 3): GENO for genotypes (19), UPheno for phenotypes (49), UBERON for anatomy (23), and MonDO for diseases (26). Prior efforts to map or integrate species-specific anatomical ontologies (24,50), for example, have been utilized in the construction of these species-neutral ontologies. The end result is a translational platform that allows a unified view of human, model and non-model organism biology.

A comparison between Monarch and existing resources is warranted. InterMine is an open-source data warehouse system used for disseminating data from large, complex biological heterogeneous data sources (51). InterMine provides sophisticated web services to support denormalized query and has been used to improve query and data access to model organism databases (52) and non-model organisms (53). InterMine is a federated approach where individual databases each can adopt and populate their own object-oriented data model, but can also align on certain aspects such as having genomic data models aligned using the SO. However, as yet genotype and phenotype modeling is not aligned, and Intermine does not provide disease matching or phenotypic search. We are currently working with InterMine to achieve harmonization in this area. Other resources, such as KaBOB (54) and Bio2RDF (38) semantically integrate various resources into large triplestores. Bio2RDF typically retains the source vocabulary of the integrated resources, whereas KaBOB is more similar to Monarch in that it maps OBO ontologies (18). Other data integration approaches include the BioThings API, exemplified by the MyVariant system (32) which aggregates variant data from multiple sources. We are currently working with the BioThings API developers to integrate these different approaches within the Dipper framework. Monarch is unique in that it aims to align both genotypic and phenotypic modeling across species and sources.

### Future directions

Future directions include bringing in phenotypic data from specialized sources and databases, incorporating a wider range of datatypes, and to extend and improve analytic methods for making cross-species inferences. Currently the core of Monarch includes primarily qualitative phenotypes described using terms from existing phenotypic vocabularies – we are starting to bring in more quantitative data, from sources such as the MPD (55) and GeneNetwork (56), in addition to expression data annotated to Uberon in BgeeDb (57). We are also extending our phenotypic search methods to incorporate Phenologs, phenotypic groupings inferred on the basis of orthologous genes (58,59). Early comparisons suggest that addition of phenologs to our suite of tools to enable genotype-phenotype inquiry across species will extend our reach in a synergistic manner (60). We therefore plan to implement this type of approach into the Monarch tool suite and website. One of the most important realizations we came across in constructing the Monarch platform was the need to better represent scientific evidence of genotype-phenotype associations. We are currently developing a Scientific Evidence and Provenance Information Ontology (SEPIO) (61) in collaboration with the Evidence and Conclusion Ontology consortium (21) and ClinGen (62) in order to classify associations as complementary, confirmatory, or contradictory. SEPIO will also integrate biological assays from the Ontology of Biomedical Investigations (63). Monarch has also been collaborating with the US National Cancer Institute’s Thesaurus (NCIT) team to integrate cancer phenotypes. Finally, Monarch has been working in the context of the Global Alliance for Genomics and Health (GA4GH) to develop a formal phenotype exchange format (<u>www.phenopackets.org</u>) that can aid phenotypic data sharing in numerous contexts such as clinical, model organism research, biodiversity, veterinary, and evolutionary biology.

## Glossary of Acronyms

| Acronym | Name | URL | Ref | PMID | NAR DB |
| --- | --- | --- | --- | --- | --- |
| Bgee | BgeeDb | http://bgee.org/ | 57 |  |  |
| BioGrid | Biological General Repository for Interaction Datasets. | https://thebiogrid.org/ | 33 | PMID:25428363 | D470-D478 |
| CL | Cell Ontology | http://obofoundry.org/ontology/cl.html | 64 | PMID:27377652 | 42/D1/D950 |
| ClinVar | ClinVar | https://www.ncbi.nlm.nih.gov/clinvar/ | 2 | PMID:26582918 | D862-8 |
| CTD | Clinical Toxicology Database | http://ctdbase.org/ | 4 | PMID:27651457 | D914-20 |
| ECO | Evidence and Conclusions Ontology | http://obofoundry.org/ontology/eco.html | 21 | PMID:25052702 |  |
| ExAC | Exome Aggregation Consortium | http://exac.broadinstitute.org/ | 6 | PMID:27535533 |  |
| FlyBase | FlyBase | http://flybase.org | 65 | PMID:26467478 | D786-92 |
| GeneNetwork | Gene Network | http://genenetwork.org | 56 |  |  |
| GENO | Genotype Ontology | https://github.com/monarch-initiative/GENO-ontology/ | 19 |  |  |
| GO | Gene Ontology | http://geneontology.org | 37 | PMID:25428369 | D1049-56 |
| GWAS | GWAS Catalog | https://www.ebi.ac.uk/gwas/ | 5 | PMID:24316577 | D1001-6 |
| HP | Human Phenotype Ontology | http://human-phenotype-ontology.org/ | 30 | PMID:24217912 | D966-D974 |
| KEGG | Kyoto Encyclopedia of Genes and Genomes | http://www.kegg.jp/ | 31 | PMID:24214961 | D199-205 |
| MGI | Mouse Genome Informatics | http://www.informatics.jax.org/ | 36 | PMID:26238262 |  |
| MonDO | Monarch Merged Disease Ontology | https://github.com/monarch-initiative/monarch-disease-ontology/ | 26 |  |  |
| MP | Mammalian Phenotype Ontology | http://obofoundry.org/ontology/mp.html | 20 | PMID:26238262 |  |
| MPD | Mouse phenome database | http://phenome.jax.org/ | 55 | PMID:24243846 | D825-34 |
| MyGene | MyGene | http://mygene.info | 32 | PMID:27154141 |  |
| OMIA | Online Mendelian Inheritance in Animals | http://omia.angis.org.au/home/ | 41 |  |  |
| OMIM | Online Mendelian Inheritance in Man | http://omim.org | 1 | PMID:25428349 | D789-98 |
| OrphaNet | Portal for rare diseases and orphan drugs | http://www.orpha.net | 3 | PMID:22422702 |  |
| Panther | PantherDB | http://pantherdb.org | 34 | PMID:26578592 | D336-42 |
| RO | Relation Ontology | http://obofoundry.org/ontology/ro.html | 17 | PMID: 15892874 |  |
| SEPIO | Scientific Evidence and Provenance Information Ontology | https://github.com/monarch-initiative/SEPIO-ontology/ | 61 |  |  |
| SO | Sequence Ontology | http://www.sequenceontology.org/ | 27 | PMID:26229585 |  |
| Uberon | Uber-anatomy ontology | http://uberon.org | 23 | PMID:25009735 |  |
| Upheno | Unified Phenotype Ontology | https://github.com/obophenotype/upheno/ | 49 | PMID:24358873 |  |
| WormBase | WormBase | http://wormbase.org | 66 | PMID:26578572 | D774-80 |
| ZFIN | Zebrafish Information Resource | http://zfin.org | 35 | PMID:26097180 |  |

## Acknowledgments

We thank members of the Undiagnosed Disease Program, the International Mouse Phenotyping Consortium, the NCI Semantic Infrastructure team, and NIF/SciCrunch for their contributions.

